# Structural basis for the unexpected activity of rifamycin B against rifampicin-resistant RNA polymerase

**DOI:** 10.64898/2026.07.13.738162

**Authors:** Hamed Mosaei, Yeonoh Shin, Valery N. Kozhevnikov, Paul G. Waddell, Michael John Hall, Katsuhiko S. Murakami, Nikolay Zenkin

**Affiliations:** Centre for Bacterial Cell Biology, Biosciences Institute, Faculty of Medical Sciences, Newcastle University, Baddiley-Clark Building, Richardson Road, Newcastle Upon Tyne, NE2 4AX, UK; Department of Biochemistry and Molecular Biology, Penn State University, University Park, PA 16802, USA; Huck Institutes of the Life Sciences, Center for Structural Biology, Penn State University, University Park, PA 16802, USA; Huck Institutes of the Life Sciences, Center for RNA Molecular Biology, Penn State University, University Park, PA 16802, USA; Hansoh Bio, LLC, 9900 Medical Center Drive, Suite 200, Rockville, MD 20850, USA; School of Geography and Natural Sciences, Northumbria University, Newcastle Upon Tyne, Tyne and Wear, NE1 8ST, UK; School of Natural and Environmental Sciences, Newcastle University, Newcastle-upon-Tyne, NE1 7RU, UK

## Abstract

Rifamycins inhibit bacterial transcription by targeting RNA polymerase (RNAP), but their clinical effectiveness is limited by the rapid emergence of resistance caused by mutations within the rifamycin-binding pocket. Rifamycin B (Rif B), one of the earliest discovered members of this antibiotic family and a precursor of clinically used derivatives, has remained poorly characterized because of its chemical instability and relatively weak antibacterial activity. Here, we revisit Rif B using biochemical and structural approaches. We show that Rif B remains sufficiently stable under assay conditions and retains inhibitory activity against RNAP variants carrying clinically relevant rifampicin-resistance mutations. We report the first crystal structure of Rif B and determine the structure of Rif B bound to bacterial RNAP. The structures reveal that the distinctive C-4 *O*-carboxymethyl substituent of Rif B forms an intramolecular interaction in the free molecule but establishes a salt bridge with fork loop 2 of the RNAP β-subunit upon binding. This additional interaction explains the reduced sensitivity of Rif B to resistance-associated substitutions and identifies the C-4 position as an underexplored site for rational rifamycin modification. These findings redefine Rif B as a mechanistically distinct rifamycin scaffold and provide new insights for developing inhibitors targeting rifampicin-resistant RNAP.

## Introduction

Rifamycins are a class of antibiotics that inhibit bacterial RNA polymerase (RNAP), the enzyme responsible for RNA synthesis (1) . Since their discovery (2, 3), rifamycins have played a central role in antimicrobial therapy, most notably in the treatment of tuberculosis (TB), an infectious disease caused by *Mycobacterium tuberculosis* and a leading cause of death worldwide. Semisynthetic derivatives such as rifampicin (Rif), rifabutin, and rifapentine remain cornerstone components of multidrug regimens against *M. tuberculosis* and are also used against a range of other bacterial pathogens (4, 5)

Rifamycins act by binding to the β-subunit of bacterial RNAP near the catalytic centre, physically obstructing the path of the nascent RNA transcript. This steric blockade prevents RNA chain extension beyond a few nucleotides, effectively halting transcription initiation. Because the Rif-binding pocket is highly conserved among bacterial RNAPs, rifamycins inhibit transcription across diverse bacterial species, contributing to their broad antibacterial activity. However, their clinical utility has been undermined by the widespread emergence of resistance (1).

In *M. tuberculosis* and many other bacteria, the primary mechanism of rifampicin resistance involves point mutations in the rpoB gene, which encodes the RNAP β-subunit, reducing drug-binding affinity (1). Additional mechanisms—including efflux-mediated changes in intracellular drug accumulation(6) and enzymatic inactivation (7)—have also been described, although mutations affecting the RNAP target remain the dominant mechanism associated with clinical rifampicin resistance in *M. tuberculosis*. Despite these challenges, rifamycins remain indispensable, and efforts continue to develop improved derivatives (8)

Among the earliest rifamycins discovered, rifamycin B (Rif B) was the first member of the family stable enough to be purified from fermentation of *Amycolatopsis mediterranei* (9). Rif B is known to undergo chemical transformation in aqueous oxygenated solutions, including reversible oxidation to Rif O, hydrolysis to Rif S, and reduction to Rif SV (Figure 1A) (10, 11). Rif B exhibits relatively weak antimicrobial activity, which has been attributed, at least in part, to limited penetration of bacterial envelopes, potentially influenced by its unique C-4 carboxylic acid-containing substituent (Figure 1A). Early efforts to modify the C-4 group produced analogues with improved antibacterial activity, but these studies were not pursued further as attention shifted toward Rif SV-derived compounds with improved stability, potency, and pharmacokinetic properties (1). Because these C-4-modified derivatives were developed before the molecular basis of rifampicin resistance was established, their activity against clinically relevant rifampicin-resistant RNAP variants and bacterial strains remains largely unexplored.

**Figure 1.**
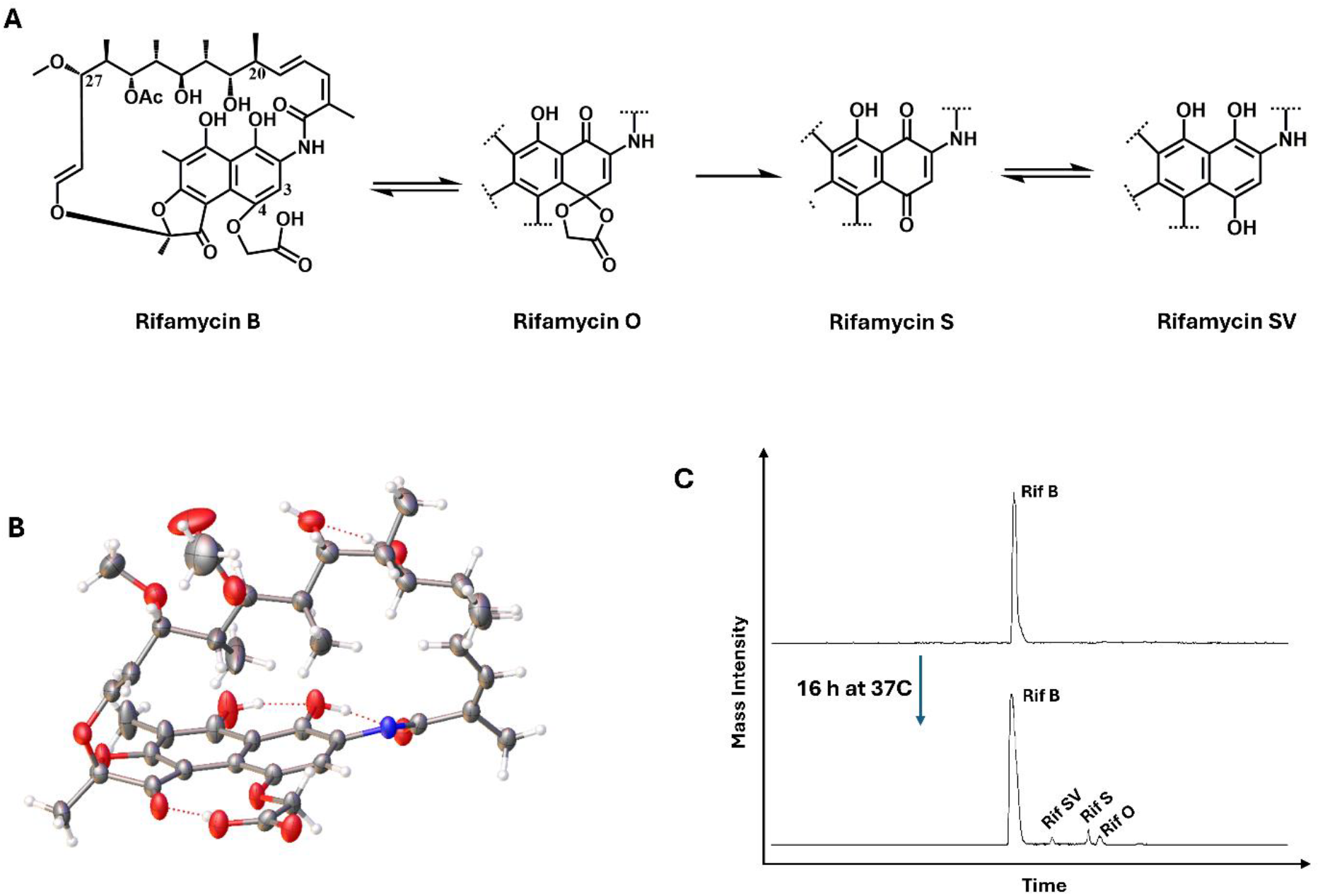
Structure and chemical conversion of rifamycin B. (A) Rifamycin B is converted to rifamycin O, followed by hydrolysis to rifamycin S, which can be reduced to rifamycin SV. (B) Crystal structure of Rif B. One crystallographically independent molecule is shown for clarity, with anisotropic displacement parameters displayed at 50% probability. Atoms are colored by element: carbon, gray; oxygen, red; nitrogen, blue; and hydrogen, white. (C) LC–MS analysis of Rif B conversion to other rifamycin derivatives over 16 h in conditions used in biochemical assays. Detected masses were as follows: [Rifamycin S + H]^+^ = 696.3006, [Rifamycin SV + H]^+^ = 698.3087, [Rif B + H]^+^ = 756.32, [Rif B + Na]^+^ = 778.3050, and [Rifamycin O + H]^+^ = 754.3013.

Rif SV, a more stable and potent downstream rifamycin lacking the C-4 substituent of Rif B, subsequently became the scaffold for clinically relevant derivatives, including rifampicin and rifapentine, which are modified at C-3, and rifaximin and rifabutin, which are modified at C-3/C-4 (12). Consequently, Rif B itself, despite its central position as a precursor of other rifamycins, has remained understudied, and definitive structural data for the unmodified molecule have not been available. This knowledge gap is particularly notable because the C-4 carboxyl-containing substituent is a distinctive feature of Rif B, yet its contribution to RNAP inhibition and sensitivity to rifampicin-resistance mutations has remained unknown(12). This raises the question of whether the C-4 carboxyl substituent of Rif B represents only a transient biosynthetic feature lost during conversion to downstream rifamycins, or whether it provides a previously unrecognized functional contribution.

Here, we show that Rif B retains activity against clinically relevant rifampicin-resistant RNAP variants. We report the first crystal structure of Rif B and determine the structure of Rif B bound to bacterial RNAP, revealing the molecular basis of its interactions with the Rif-binding pocket that lead to such activity. Our findings redefine Rif B as a mechanistically distinct rifamycin scaffold and reveal how modification at C-4 influences RNAP inhibition and resistance sensitivity.

## Results

### Structural and chemical characterization of Rifamycin B

Although crystal structures of chemically modified Rif B derivatives have been reported(13, 14), an atomic three-dimensional structure of the unmodified molecule has not been described. We therefore determined the single-crystal X-ray structure of Rif B to define its conformation and provide a structural basis for comparison with the RNAP-bound state (Figure 1B). Single crystals of Rif B suitable for single crystal X-ray analysis were obtained from the slow cooling of an acetonitrile solution. The structure was observed to be an acetonitrile solvate with two crystallographically independent molecules in the asymmetric unit and *P*2_1_2_1_2_1_ space group symmetry (Table S1). The crystal structure confirms the presence of an *O*-carboxymethyl substituent at the 4-position of the 1,4-dihydroxynaphthalene ring system. In the crystal structure the 4-*O*-carboxymethyl group is stabilized through an intramolecular H-bond with the C-11 carbonyl, forming a ten-membered H-bonded ring structure, classified as an S(10) hydrogen bonding motif using Etter’s graph set notation (15). This structure provides the first complete crystallographic analysis of Rif B, unambiguously validates its structure and absolute stereochemistry, and establishes a reference model for comparison with Rif B bound to *Thermus thermophilus* RNAP.

Because Rif B is known to undergo chemical conversion in aqueous oxygenated solutions, we next examined its stability under conditions relevant to the biochemical and antibacterial assays. LC–MS analysis showed gradual conversion of Rif B into Rif O, Rif S, and Rif SV over time (Figure 1C). However, conversion was very slow: after 16 h at 37°C, Rif B remained the predominant species. Thus, under the conditions used in the activity assays, Rif B is sufficiently stable for the observed biological effects to be attributed to it, rather than to downstream transformation products.

### Rifamycin B inhibits Rifampicin-resistant RNA polymerases

We first tested whether Rif B inhibits bacterial RNAP through the canonical rifamycin mechanism. In vitro transcription assays were performed with wild-type *E. coli* RNAP on a linear DNA template containing the strong T7A1 promoter (Figure 2A). In the absence of inhibitor, RNAP produced full-length run-off and terminated transcripts. Addition of Rif B reduced formation of full-length RNA and led to accumulation of short abortive trinucleotide and tetranucleotide products, consistent with inhibition during early transcription initiation. This mode of inhibition results from Rif B binding in the path of the nascent RNA, preventing transcript extension and promoting release of short abortive products. The half-maximal inhibitory concentrations (IC_50_) of Rif B and Rif against wild-type *E. coli* RNAP were comparable (Figure 2B and C), consistent with a shared general mechanism of binding.

**Figure 2.**
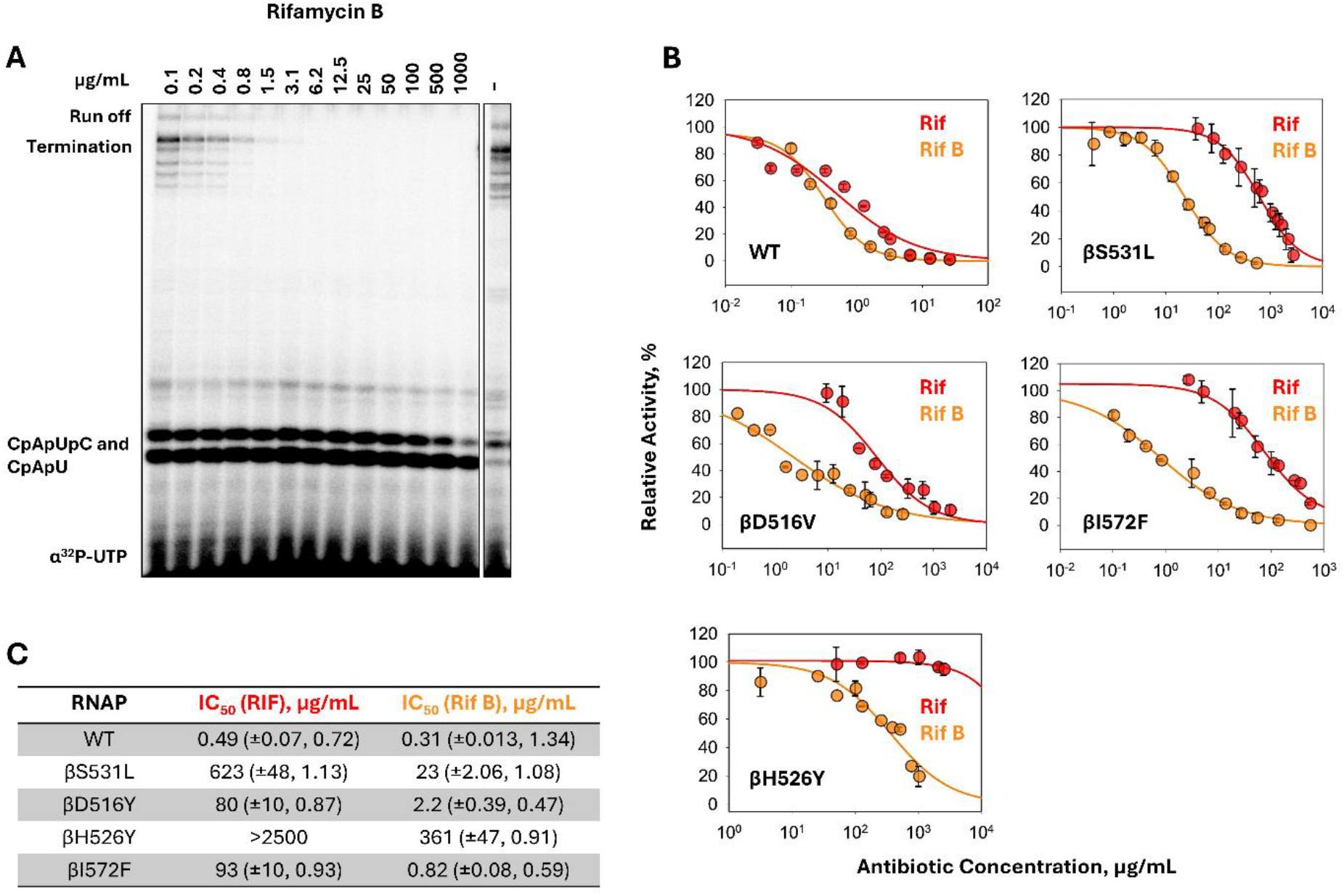
Inhibition of *E. coli* RNAP by rifamycin B and rifampicin. (A) In vitro transcription by *E. coli* RNAP initiated with CpA on a linear DNA template containing the T7A1 promoter, in the presence of increasing concentrations of Rif B (0–1000 µg/mL). Run-off, termination, and abortive tri-and tetranucleotide products are indicated. (B) Dose-dependent inhibition of wild-type and Rif-resistant *E. coli* RNAP variants by Rif and Rif B. Error bars represent ± SD. (C) IC_50_ values calculated from the inhibition curves shown in panel B. Values in parentheses indicate ± SE and Hill slope.

We next tested whether Rif B retained activity against RNAP variants carrying substitutions associated with rifampicin resistance. Purified *E. coli* RNAPs containing βS531L, βD516Y, βH526Y, or βI572F substitutions were assayed under the same conditions. These mutations correspond to those most frequently found in *M. tuberculosis*, with alterations at βS531, βD516, and βH526 (*E. coli* numbering) collectively accounting for over 70% of Rif-resistant clinical isolates (1). Compared with Rif, Rif B retained substantially greater inhibitory activity against all rifampicin-resistant RNAP variants tested (Figure 2B and C). The difference was most pronounced for βS531L and βD516Y, for which Rif B was approximately 30-fold and 40-fold more active than Rif, respectively. Notably, while the βH526Y substitution rendered RNAP completely insensitive to Rif (no inhibition at 2.5 mg/mL), Rif B still inhibited this mutant enzyme at high concentrations.

These results show that Rif B acts as a bona fide rifamycin-class RNAP inhibitor but retains activity against RNAP variants that confer resistance to Rif. This reduced sensitivity to resistance-associated substitutions suggested that Rif B forms distinct interactions within the RNAP’s Rif-binding pocket, prompting structural analysis of the RNAP–Rif B complex.

### Structural basis of RNAP Inhibition by Rif B

To elucidate the structural basis for the enhanced activity of Rif B relative to Rif against Rif-resistant RNAP variants, we determined the X-ray crystal structure of Rif B in complex with the *Thermus thermophilus* (Tth) RNAP σ^A^ holoenzyme assembled on a *pyrG* promoter-containing DNA scaffold (Table S2)(8). The electron density map revealed unambiguous density for Rif B within the Rif-binding pocket (Figure 3A). The overall binding mode of Rif B is similar to that of Rif, and the Rif B molecule could be modeled into the electron density without requiring major conformational adjustments of either the ligand or the surrounding protein. The Rif B binding is stabilized through a network of interactions with Rif-binding pocket residues, including Q510, Q513, S531, R540 (side chain; *E. coli* numbering), and F514 (main chain; *E. coli* numbering) (Figure 3B). Notably, Rif B forms an additional salt bridge between its 4-*O*-carboxymethyl group and the R540 side chain of β-subunit fork loop 2. Thus, Rif B binds within the canonical Rif-binding pocket but establishes an additional C-4-dependent interaction not present in Rif, providing a structural basis for the biochemical differences observed with Rif-resistant RNAP variants.

**Figure 3.**
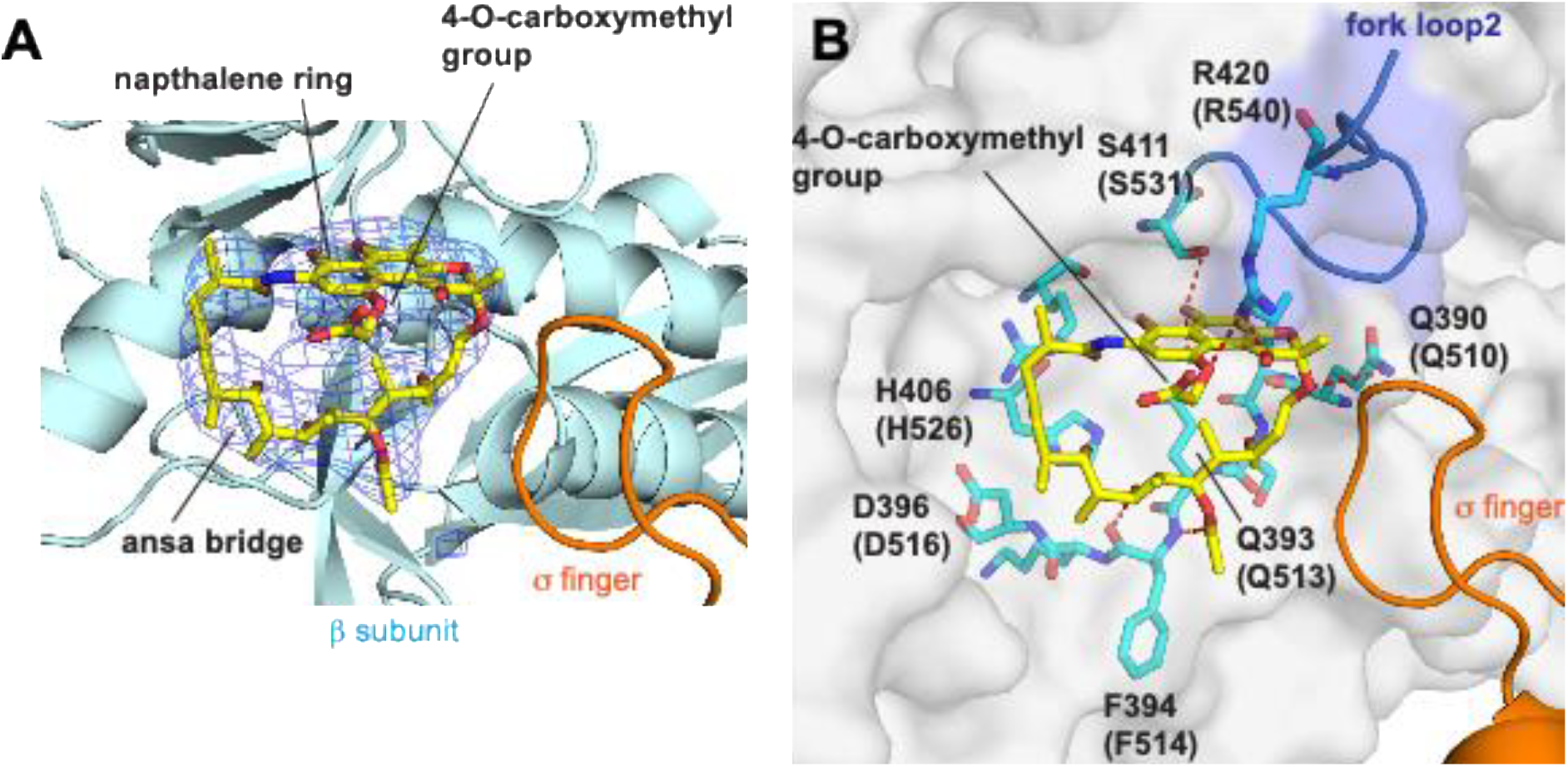
Structure of RNAP and Rif B complex. (A) Electron density map (blue mesh) of Rif B (stick model) in complex with *T. thermophilus* RNAP (β subunit: cyan, σ factor: orange). Ansa bridge, naphthalene ring and 4-*O*-carboxymethyl group of Rif B are indicated. (B) RNAP and Rif B interaction. Residues of Rif-binding pocket of β subunit (*T. thermophilus* number, corresponding residues in *E. coli* RNAP are shown in parentheses) participating in hydrogen bonds and salt bridge (red dashed lines) are shown in stick models with β subunit surface (white transparent) and σ finger (orange cartoon model). Fork loop 2 is colored in blue. Amino acid residues involved in interaction with Rif B are shown as stick models in cyan.

## Discussion

Rifamycins remain among the most important inhibitors of bacterial transcription and key components of first-line treatment for tuberculosis. However, their clinical use is limited by the rapid emergence of resistance, most commonly through mutations in the RNAP Rif-binding pocket. Therefore, it is important to understand how natural rifamycin variants inhibit RNAP and whether alternative chemical features can improve activity against resistant enzymes. Rif B occupies an unusual position in this context: it was among the earliest rifamycins isolated, but its molecular properties have remained poorly defined. The principal finding of this work is that Rif B retains inhibitory activity against RNAPs carrying mutations that confer resistance to other rifamycins derived from this scaffold.

The first crystal structure of unmodified Rif B, together with its complex with RNAP, provide a structural framework for understanding this activity. In the apo structure, the C-4 *O*-carboxymethyl group forms an intramolecular hydrogen bond with the C-11 carbonyl, whereas in the RNAP-bound state this interaction is replaced by a salt bridge between the C-4 *O*-carboxymethyl group and βR540 in fork loop 2 of the Rif-binding pocket. Thus, a substituent historically associated with reduced antibacterial activity directly contributes to binding to RNAP.

The location of this interaction provides a structural explanation for Rif B activity against Rif-resistant RNAP variants, particularly the S531L mutant. Although the S531L substitution does not significantly alter the overall shape of the Rif-binding pocket, previous structural work showed that the deeper insertion of the naphthyl ring in Rif-resistant variants forces fork loop 2 open, exposing the ring to solvent (16). We propose that the interaction between βR540 and the C-4 carboxymethyl group stabilizes fork loop 2 and prevents this opening during Rif B binding. Thus, a chemical feature removed during rifamycin optimization may retain mechanistic value by providing an additional interaction within the target.

The orientation of the C-4 substituent also highlights opportunities for future rifamycin design. The carboxymethyl group faces σ region 3.2 (the σ finger), which contains acidic residues in most primary bacterial σ factors. Because both the Rif B carboxyl group and the σ finger are negatively charged, no direct interaction occurs in the current structure. Modifications at the C-4 position introducing polar or positively charged groups may therefore create additional interactions with σ region 3.2. Combining such modifications with other rifamycin extensions that expand interactions within RNAP, such as those observed in kanglemycin A (8) could provide routes toward derivatives with improved activity against Rif-resistant variants.

Indeed, while several clinically used rifamycins contain modifications at C-3 or C-3/C-4 positions, these substituents do not establish equivalent interactions with fork loop 2 (17). Similar engagement of this region has previously been reported for specifically designed bulky benzoxazinorifamycin derivatives (18), which were suggested to contact the corresponding arginine residue within the Rif-binding pocket. Thus, Rif B reveals that a naturally occurring C-4 substituent can provide an additional interaction within the Rif-binding pocket, contributing to its reduced sensitivity to Rif-resistance mutations.

More broadly, Rif B should not be viewed only as an unstable precursor or degradation-prone intermediate. Its ability to form a specific interaction with RNAP suggests that Rif B may have functional properties distinct from its downstream conversion products. This is consistent with the idea that interconverting rifamycins may not simply represent inactive intermediates on the way to a final product, but chemically distinct congeners with different target interactions. Revisiting historically overlooked natural products with modern structural and biochemical tools can therefore uncover interactions that were missed during optimization campaigns focused mainly on stability, potency, and pharmacokinetics.

Together, our results identify the Rif B C-4 substituent as an underexplored site for rational rifamycin modification. Preserving the βR540 interaction, introducing polar or positively charged groups capable of engaging σ region 3.2, or combining C-4 modifications with additional interactions found in other rifamycin scaffolds may provide routes to derivatives with improved activity against Rif-resistant variants and bacterial strains.

## Materials and Methods

### LC–MS analysis of rifamycin B conversion

LC–MS analysis of rifamycin B (Rif B) was performed using an Agilent 1260 HPLC system. Rif B was dissolved in transcription buffer (20 mM Tris-HCl, pH 7.9; 40 mM KCl; 10 mM MgCl_2_) and incubated at 37°C for 16 h. Samples (1 μL) were injected onto a Raptor ARC-18 column (150 × 2.1 mm; Restek) operated at a flow rate of 0.2 mL/min. Compounds were separated using a 30-min linear gradient from 5% to 100% acetonitrile, with the mobile phase supplemented with 0.1% formic acid. Mass spectra were acquired in positive-ion mode using a Bruker MicrOTOF II time-of-flight mass spectrometer.

### Purification of *E. coli* RNAP core enzyme and σ70 for transcription assays

Rifampicin-resistant mutations were introduced into the pETLRpoB plasmid, which encodes the *E. coli* RNAP β subunit with an N-terminal 6×His tag (19), by site-directed mutagenesis using the QuikChange II kit (Stratagene), as previously described(8). Wild-type and mutant pETLRpoB plasmids were co-transformed with pACYCDuet-1_Ec_rpoZ (encoding the ω subunit) into *E. coli* T7 Express cells (New England Biolabs).

Protein expression and purification of core RNAP were carried out as described previously(8). The N-terminally 6×His-tagged *E. coli* σ^70^ subunit was purified separately following established procedures (20). Purified RNAP core and σ^70^ subunits were quantified spectrophotometrically and stored at −20°C in storage buffer (20 mM Tris-HCl, pH 7.9; 50 mM NaCl; 50% glycerol) until use.

### In vitro transcription

In vitro transcription assays were performed as described previously(8). Briefly, 1 pmol of wild-type or mutant *Escherichia coli* RNA polymerase (RNAP) core enzyme was mixed with 3 pmol of σ^70^ and 0.1 pmol of a linear DNA template carrying the *T7A1* promoter in transcription buffer (20 mM Tris-HCl, pH 7.9; 40 mM KCl; 10 mM MgCl_2_) in a total volume of 7 μl. Rif B (Merck) was added in 1 μl of DMSO (ranging from 1 µg/mL to 10 mg/mL) or an equivalent volume of DMSO for control reactions. Transcription was initiated by adding 2 μl of nucleotide mixture in transcription buffer containing, at final concentrations, 25 μM CpA, 100 μM ATP, CTP, and GTP, and 10 μM [α-^32^P]UTP (20 Ci/mmol; Hartmann Analytic). Reactions were incubated for 10 min at 37°C and stopped by the addition of formamide-containing loading buffer.

RNA products were resolved on 20% (8 M urea) polyacrylamide gels, visualized using a PhosphorImager (Cytiva), and quantified with ImageQuant software (Cytiva). Transcriptional activity (sum of run-off and termination products) was normalized to the antibiotic-free control, defined as 100%. Data represent means ± standard deviations from at least three independent experiments. Dose– response curves were fitted using a four-parameter logistic equation by nonlinear regression (SigmaPlot; Systat Software). For direct comparison, IC_50_ values for RIF were extracted from the previously published dataset(8) and plotted alongside newly obtained Rif B data generated under identical assay conditions.

### Crystallization of RNAP-Rif B complexes

*Thermus thermophilus* RNAP σ^A^ holoenzyme in complex with the *pyrG* promoter DNA was prepared and crystallized as described earlier(21). For RNAP-Rif B complex preparation, RNAP crystals were transferred to the corresponding cryoprotectant solutions supplemented with 1 mM Rif B and incubated at 22 °C overnight. Following soaking, crystals were rapidly cryo-cooled by immersion in liquid nitrogen prior to data collection.

X-ray diffraction data were collected at the Macromolecular Diffraction (MacCHESS) F1 beamline at the Cornell High Energy Synchrotron Source (Cornell University, Ithaca, NY). The datasets were processed using HKL2000 (22). Initial phases were obtained by molecular replacement with a previously reported structure of the *T. thermophilus* RNAP–*pyrG* promoter complex(21) used as search models. Structural refinement was carried out using the Phenix software suite (23), employing iterative rigid-body and positional refinement and reference-model restraints. Electron density maps revealed well-defined density corresponding to Rif B, which was absent from the initial models; the ligand was subsequently built into the maps using Coot (Emsley and Cowtan, 2004).

Data collection and refinement statistic for RNAP–Rif B complex is summarized in Table S2. Final coordinate and structure factor have been deposited in the Protein Data Bank under the accession codes listed in Table S2.

### Small molecule X-ray crystallography of Rif B

Rif B was dissolved in acetonitrile at 65 °C to a final concentration of 5 mg/mL. Slow cooling of the solution down to room temperature resulted in formation of crystals suitable for single-crystal X-ray analysis.

Crystal structure data were collected on an XtaLAB Synergy HyPix-Arc 100 diffractometer using copper radiation (λ_CuKα_ = 1.54184 Å). Data were collected at 150 K using an Oxford Cryosystems CryostreamPlus open-flow N_2_ cooling device. Intensities were corrected for absorption using a multifaceted crystal model created by indexing the faces of the crystal for which data were collected (24). Cell refinement, data collection and data reduction were undertaken via the software CrysAlisPro (25)

All structures were solved using XT (26) and refined by XL (27) using the Olex2 interface (28). All non-hydrogen atoms were refined anisotropically and hydrogen atoms were positioned with idealised geometry, with the exception of those bound to heteroatoms, the positions of which were located using peaks in the Fourier difference map. The displacement parameters of the hydrogen atoms were constrained using a riding model with U_(H)_ set to be an appropriate multiple of the U_eq_ value of the parent atom.

Part of one of the molecules comprising the main residue has been modelled as disordered over two positions. The occupancies of the two parts were refined independently of the atomic displacement parameters. The geometry was restrained using the SADI card and the displacement parameters of all partially-occupied non-hydrogen atoms were restrained using the SIMU card. The structure contains solvent accessible voids. These voids appear to contain ca. 3.5 acetonitrile molecules disordered over multiple positions. As no sensible model for the solvent was forthcoming the associated electron density was treated using the Olex2 solvent mask routine.

## Acknowledgments

We thank the staff at the MacCHESS for support with crystallographic data collection for the RNAP-Rif B complex, and Prof. Michael Probert (Newcastle University) for crystallization support.

## Accession Codes

CCDC 2570401 contains the supplementary crystallographic data for the single crystal structure of Rif B determined in this paper. These data can be obtained free of charge via www.ccdc.cam.ac.uk/data_request/cif, or by emailing, or by contacting The Cambridge Crystallographic Data Centre, 12 Union Road, Cambridge CB2 1EZ, UK; fax: +44 1223 336033. The atomic coordinates and structure factors for *T. thermophilus* RNAP – Rif B complex have been deposited in the Protein Data Bank (PDB: 12YY).

