## Supplementary material for "Structural basis for the unexpected activity of rifamycin B against rifampicin-resistant RNA polymerase"

**Table S1: Crystal data and structure refinement for Rif B.**

| | |
|---|---|
| CCDC Number | 2570401 |
| Empirical formula | $C_{80}H_{101}N_3O_{28}$ |
| Formula weight | 1552.63 |
| Temperature/K | 150.0(2) |
| Crystal system | orthorhombic |
| Space group | $P2_12_12_1$ |
| a/Å | 16.0358(3) |
| b/Å | 21.0354(5) |
| c/Å | 26.5588(5) |
| α/° | 90 |
| β/° | 90 |
| γ/° | 90 |
| Volume/Å$^3$ | 8958.8(3) |
| Z | 4 |
| $\rho_{calc}$g/cm$^3$ | 1.151 |
| μ/mm$^{-1}$ | 0.727 |
| F(000) | 3304.0 |
| Crystal size/mm$^3$ | 0.18 × 0.02 × 0.02 |
| Radiation | CuKα (λ = 1.54184) |
| 2Θ range for data collection/° | 5.36 to 153.83 |
| Index ranges | -9 ≤ h ≤ 19, -24 ≤ k ≤ 25, -33 ≤ l ≤ 33 |
| Reflections collected | 53662 |
| Independent reflections | 17561 [$R_{int}$ = 0.0643, $R_{sigma}$ = 0.0688] |
| Data/restraints/parameters | 17561/1057/1093 |
| Goodness-of-fit on F$^2$ | 0.987 |
| Final R indexes [I>=2σ (I)] | $R_1$ = 0.0478, w$R_2$ = 0.1064 |
| Final R indexes [all data] | $R_1$ = 0.0779, w$R_2$ = 0.1192 |
| Largest diff. peak/hole / e Å$^{-3}$ | 0.30/-0.22 |
| Flack parameter | -0.05(8) |

**Table S2. Collection of crystallographic data and refinement statistics of the *Tth* RNAP-*pyr*G1 promoter complex with RifB (PDB code: 12YY)**

| | |
|---|---|
| Data collection | |
| Space group | C2 |
| Cell dimensions | |
| a (Å) | 185.460 |
| b (Å) | 101.256 |
| c (Å) | 294.346 |
| β (°) | 98.748 |
| Resolution (Å) | 50 – 2.90 (2.95-2.90) |
| Total reflections | 369,465 |
| Unique reflections | 114,412 |
| Redundancy | 3.2 (2.2) |

| | |
|---|---|
| Completeness (%)* | 95.6 (63.9) |
| I / σ* | 16.2 (1.2) |
| Rsym (%)* | 12.3 (>112) |
| CC1/2* | (0.445) |
| Refinement | |
| Resolution (Å) | 43.72 – 3.00 (3.08-3.00) |
| Rwork | 0.205 (0.352) |
| Rfree | 0.251 (0.379) |
| No. of atoms | 28,486 |
| macromolecules | 28,427 |
| ligand | 102 |
| solvent | 0 |
| R.m.s deviations | |
| Bond length (Å) | 0.011 |
| Bond angles (°) | 1.53 |
| Clashscore | 10.27 |
| Ramachandran favored, % | 96.92 |
| Ramachandran outliers, % | 0.40 |
| Rotamer outliers, % | 0.20 |
| Average B-factor | 92.73 |
| macromolecules | 92.13 |
| ligand | 70.89 |

*Highest resolution shells are shown in parentheses
